# Nipah virus Malaysia and Bangladesh strain-induced pathogenesis in mice lacking type I interferon receptor signaling

**DOI:** 10.64898/2025.12.24.696352

**Authors:** Lucia Amurri, Olivier Reynard, Ilona Ronco, Daniel Déri, Bernadett Pályi, Julia Spanier, Jennifer Skerra, Olivia Terceve, Thibaut Larcher, Ulrich Kalinke, Zoltán Kis, Branka Horvat, Mathieu Iampietro

**Affiliations:** CIRI, Centre International de Recherche en Infectiologie, INSERM U1111, CNRS, UMR5308, Univ Lyon, Université Claude Bernard Lyon 1, École Normale Supérieure de Lyon, 21 Avenue Tony Garnier, 69007 Lyon, France; Institute for Experimental Infection Research, TWINCORE, Centre for Experimental and Clinical Infection research, a joint venture between the Helmholtz-Centre for Infection Research and the Hannover Medical School, Hannover, Germany; National Biosafety Laboratory, National Center for Public Health and Pharmacy, Budapest, Hungary; Semmelweis University, Institute of Medical Microbiology, Budapest, Hungary; European Research Infrastructure on Highly Pathogenic Agents (ERINHA-AISBL), Brussels, Belgium; UMR703, PAnTher, APEX, Oniris VetAgroBio, La Chantrerie, CS 40706, 44307 Nantes, France; Cluster of Excellence - Resolving Infection Susceptibility (RESIST, Hannover Medical School, EXC 2155), Hannover, Germany

## Abstract

Nipah virus (NiV) is a zoonotic highly pathogenic *Paramyxovirus* inducing lethal outbreaks of encephalitis and severe acute respiratory syndrome (SARS) with an average case-fatality rate of 75%. Two viral strains, NiV-Malaysia (NiV-Mal) and NiV-Bangladesh (NiV-Ban), associated to distinct geographical distribution, route of transmission, symptoms and lethality have been described. Due to the permanent threat of these emerging infections and the lack of approved therapeutics, it is crucial to improve our understanding regarding NiV-associated pathogenesis. Mice represent a small and accessible animal model, provided with numerous biological tools for the functional assessment of different genes related to antiviral response. Here, we analyze the susceptibility of mice deficient for type I interferon receptor (IFNAR KO) to infection with either NiV-Mal or NiV-Ban through intraperitoneal or intranasal routes. Our results show that IFNAR KO mice are susceptible to NiV-Ban infection via intraperitoneal route, although to a lesser extent than NiV-Mal, and develop encephalitis and pulmonary syndrome with viral propagation to different organs and lethal outcome in 60% of infected animals. In addition, intranasal administration of both viral strains led to a subclinical infection with viral replication in brain and lungs and production of virus-specific neutralizing antibodies. These results indicate IFNAR KO mice as a small animal model permitting comparative studies of the immunopathogenesis perpetrated by both NiV-Mal and NiV-Ban infections.

**Author summary:** Availability of small animal models represents a major issue to characterize virus-associated pathogenesis to further implement therapeutic strategies. Previous studies performed with NiV-Mal and NiV-Ban strains in wild-type (WT) mice did not indicate signs of NiV disease. However, when challenged with NiV-Mal, IFNAR KO mice suffered fatal outcomes, thus providing useful information on viral immunopathogenesis. Surprisingly, while being the more frequently re-emerging virus strain, no infection studies with NiV-Ban have been performed in IFNAR KO mice so far. Here, we sought comparing pathogenesis after NiV-Mal- and NiV-Ban-infection following IP and IN inoculations in IFNAR KO mice. Indeed, our results highlighted that contrary to fatal IP challenge, IN inoculation lead to a subclinical infection with both NiV strains. Moreover, we determined that NiV-Mal is more pathogenic than NiV-Ban following IP infection. Finally, distinct histopathological manifestations and tropism were associated with each viral strain and specific route of infection. Overall, our study implies that IFNAR KO mice represent a useful animal model to study strain-specific NiV pathophysiology.

## Introduction

Nipah virus (NiV) is an emerging zoonotic *Paramyxovirus* causing highly lethal outbreaks in South-East Asia, Bangladesh and India (1). NiV is naturally hosted by *Pteropus* fruit bats, which are widespread in tropical and subtropical areas of Asia and Indian/Pacific Ocean islands and Australia (2,3). Due to its high pathogenicity, its human-to-human transmission and the lack of approved treatments or vaccines, World Health Organization has included NiV in the Blueprint list of epidemic threats requiring urgent research and development actions (4). Moreover, climate change, deforestation and intensive farming are progressively disrupting the habitats of reservoir animals and increasing animal-human contacts, leading to an increased risk for epidemic or pandemic (5–9).

Two NiV strains have been described, NiV Malaysia (NiV-Mal) and NiV Bangladesh (NiV-Ban), that share 91.8% of nucleotide sequence homology but show remarkable differences in epidemiology and human pathology (8,10). The first NiV outbreak was registered in Malaysia in 1998-1999, where the virus was first transmitted from asymptomatic fruit bats to pigs as intermediate hosts before infecting humans (11). This led to 265 cases of acute encephalitis and 105 deaths (case fatality rate, CFR=40%), mostly among pig farmers who were exposed to infected animals.

A second NiV strain emerged in Bangladesh in 2001 through a direct spillover from bats to humans following the consumption of contaminated date palm sap (12). Indeed, in this geographical area sap is harvested from palm trees using reservoirs where bats can approach and contaminate the juice with saliva and/or urine (13). Moreover, in addition to animal-to-human transmission, respiratory human-to-human transmission has been reported (14). While NiV-Mal is commonly associated with lethal encephalitis, NiV-Ban patients mainly display severe respiratory syndrome in addition to neurological symptoms. Since 2001, NiV-Ban is re-emerging frequently in Bangladesh and India, with an average CFR around 75%, reaching up to 100% in some outbreaks and presenting constant threat for new episodes and possible pandemic spread (15–17).

NiV can infect a wide range of mammals by interacting with the highly-conserved and ubiquitously expressed Ephrin B2 and B3 receptors (18–20). However, while humans, pigs and different animal models such as non-human primates (NHP), hamsters and ferrets, develop clinical symptoms with high lethality following NiV infection, mice clear the virus despite being permissive to infection (21–23). To efficiently and better characterize viral pathogenesis, convenient animal models are needed. Indeed, while several studies using the hamster model have allowed the establishment a pre-clinical approach including both NiV strains and both intraperitoneal (IP) and intranasal (IN) route of infections (24), the development of an appropriate mouse model would represent an important progress to further decipher NiV pathogenesis *in vivo*.

In previous studies using NiV-Mal infection in both wild type (WT) and Interferon α/β receptor knock out (IFNAR KO) mice, all WT animals survive both IP and IN challenge while 100% and 60% of IFNAR KO mice succumb to NiV-Mal challenge, respectively (23,25–27). In addition, viral dissemination in the brain, lungs, spleen and liver, along brain-associated lesions, leading to neurological symptoms, were observed in IP-inoculated IFNAR KO mice (25). Afterwards, additional works have been conducted on NiV infection in WT mice (28,29), notably Dups *et al.* compared NiV-Mal and NiV-Ban infections in intranasally infected WT mice (28). All animals survived without developing disease, and both viral RNA and viral antigens were detected in the lungs of infected animals only, suggesting that WT mice were able to control viral dissemination, with no significant differences between both viral strains. Moreover, neutralizing antibodies (nAb) response was not detected. These observations were recently confirmed by Edwards *et al.*, who analyzed NiV-Ban IN infection in WT mice where all animals survived without displaying neither clinical symptoms nor histological lesions, with viral detection only in the lungs and a mild production of nAb in 3 out of 5 mice (29).

Importantly, contrary to NiV-Mal, NiV-Ban infection has not been evaluated in IFNAR KO mice yet. Here, we compare pathogenesis in both WT and IFNAR KO mice infected with NiV-Mal or NiV-Ban through IP or IN route.

First, we demonstrated the permissiveness of primary murine embryonic fibroblasts (MEFs) derived from WT and IFNAR KO mice to both NiV strains *in vitro*. Then, we analyzed the susceptibility of both mouse genotypes to NiV-Mal or NiV-Ban following IP or IN infection *in vivo*. We observed that NiV-Mal was more virulent than NiV-Ban in IP-infected IFNAR KO mice. Surprisingly, neither NiV-Mal nor NiV-Ban lead to fatal outcome following IN infection in IFNAR KO mice. Moreover, viral genome quantification and histopathological analysis were performed on harvested organs to determine the magnitude of viral spread and tropism. While IN virus inoculation was associated to a subclinical infection in all experimental groups, IP infection exerted significantly higher viral replication and tissular pathology in IFNAR KO compared to WT mice. Interestingly, both viral strains presented different cellular tropism and syncytia distribution in the lungs of IFNAR KO animals.

Globally, these results suggest that IFNAR KO mice can be used as a small animal model for fundamental and pathophysiology studies of both NiV-Mal and NiV-Ban infections, taking advantage of numerous biological tools available for murine studies.

## Materials and methods

### Ethical statement

Animal experiments were performed in the biosafety level 4 (BSL-4) facility of the National Biosafety Laboratory in Budapest according to the guidelines of the European Communities Council Directive (86/609 EEC) and were approved by the Hungarian National Authority (Scientific Ethics Council for Animal Experiments, PE/EA/1456-7/2020).

### Mice

Wild type (WT) C57BL/6J mice (Charles River) and B6.129S2-Ifnar1tm1(Neo)Agt (IFNAR-KO) mice (30) were bred under specific pathogen-free conditions in the central mouse facility of the Helmholtz Centre for Infection Research (Brunswick), at TWINCORE (Centre for Experimental and Clinical Infection Research, Hanover, Germany) and at Plateau de Biologie experimentale de la souris (PBES) in Lyon. Mouse experimental work was carried out using 3-week-old to 6-week-old mice with equal male-female distribution. Mice were kept under BSL-4 conditions in individually ventilated cages (IsoRat ISO48NFEEU, Techniplast, Italy), housed on autoclaved corn bedding (Rehofix, J. Rettenmaier & Söhne GmbH + CO KG, Rosenberg, Germany), fed autoclaved food (VRFI (P), Special Diet Services) ad libitum, given osmosis-treated water provided in bottles ad libitum, and the cages were equipped for proper environmental enrichment.

### Cell lines

WT or IFNAR KO primary murine embryonic fibroblasts (MEFs) were isolated from murine embryos obtained from female pregnant mice 13 days after conception (26,31) and cultured in Dulbecco’s modified Eagle’s medium (DMEM) GlutaMAX supplemented with 10% heat-inactivated fetal bovine serum (FBS), 0.2% 2-mercaptoethanol, 1% HEPES, 1% nonessential amino acids, 1% sodium pyruvate, and 2% penicillin-streptomycin mix at 37°C with 5% CO2.

WT and IFNAR KO MEFs were plated in 12-well plates (1.5×10^5^ cells/well) or in 24-well plates (5×10^4^ cells/well) for immunofluorescence or RT-qPCR analysis, respectively. MEFs were then infected in the Jean Mérieux BSL-4 facility in Lyon with NiV-Mal or NiV-Ban at a multiplicity of infection (MOI) of 0.3 plaque-forming units (PFU)/cell (according to the viral titer previously calculated on Vero E6 cells) for 1h at 37°C in DMEM 0% FBS. Then, virus-containing medium was removed and substituted with fresh complete medium before being incubated for 24h or 48h at 37°C.

### Viruses

NiV-Mal (isolate UMMC1 GenBank-AY029767) and NiV-Ban isolate SPB200401066 (GenBank AY988601, kindly provided by the Center for Disease Control and Prevention, Atlanta USA) were prepared by infecting Vero E6 cells (ATCC CRL-1586TM) cultured in DMEM, in the INSERM Jean Mérieux BSL-4 in Lyon, France.

### Intraperitoneal and intranasal infection of mice

All work involving live NiV-Mal and NiV-Ban infectionswas performed under BSL-4 conditions. All mice were inoculated intraperitoneally with 10^6^ PFU of NIV-Mal or NiV-Ban or intranasally with 10^5^ PFU of NiV-Mal or NiV-Ban under inhalational anesthesia with isoflurane (isoflurane: ISOFLUTEK 1000 mg/g, Laboratorios Karizoo S.A.; anesthesia station: MiniHUB-V3, TEM SEGA, France).

Following infection, animals were weighed and monitored on a daily basis as part of a scoring system to follow the symptoms and determine endpoint euthanasia. After NiV-Mal infection, animals were monitored for 32 days and organs were collected at the end of the experiment or at the day of death. For NiV-Ban infection, animals were monitored during 1-month post-infection, and in addition programmed euthanasia was performed for 4 hours, 1 day, 2 days, 4 days, 7 days or 10 days post-infection (d0+4h, d1, d2, d4, d7, d10), and at the end of the protocol in case of mice which survived infection. For graphical representations of qPCR data and statistical analyses, murine samples were pooled and grouped into early (from d0+4h to d1), mid (from d2 to d7) and late (after d7 until d32) phase of infection.

### Sample collection and preparation

Euthanasia with CO_2_ was conducted under inhalational anesthesia with isoflurane, followed by blood collection via cardiac puncture and autopsy. During autopsy, brains, lungs, spleens and livers were collected for further nucleic acid isolation and histological analysis. Organs were divided for molecular testing and were collected and frozen immediately at -80°C without any medium until further preparation. Parts of organs for immunohistochemistry were collected into fixative 10 V/V% formaldehyde-PBS solution. Then, 25-30 mg of the thawed organs were taken into MagNA Lyser Green Beads tubes (Roche Diagnostics GmbH, Mannheim, Germany) containing 600 µL of 1:100 mixture of β-mercaptoethanol and Buffer RLT (QIAGEN GmbH, Hilden, Germany). After homogenization using MagNA Lyser instrument (Roche Diagnostics GmbH), samples were centrifuged for 5 minutes at 12000 rpm. Supernatants were transferred into new tubes and after 10 min of incubation at room-temperature, an equal volume of 70 V/V% ethanol was added into each tube.

### Seroneutralization

Seroneutralization assay was performed on mice euthanized after day 7. Serum samples were stored at -80°C without previous thawing and were analyzed without heat inactivation. Serum dilution and assay procedures were done according to manufacturer’s instructions under BSL-4 conditions. Sera were diluted in 7-point serial two-fold dilutions in triplicate in serum-free DMEM in sterile 96-well microtiter plates. Positive and negative control sera were also applied, the latter with and without serum as well. Equal volume of 60±20 TCID50 NiV-Mal or NiV-Ban was added into each well and incubated for 1 h at 37°C. Then, samples were transferred on a Vero E6 cells monolayer (approximately 1,8×10^5^ cells/mL) maintained in serum-free DMEM in 96-well plates (TPP, Switzerland). The neutralizing antibody titer of each sample was determined by the lack of cytopathic effect (CPE) after a 5 days incubation at 37°C. For triplicates the geometric mean of end dilutions was calculated and reported as a neutralizing antibody titer (nAb).

### RNA extraction and RT-qPCR

For murine samples, total nucleic acid extraction was performed using QIAsymphony SP instrument with QIAsymphony DSP Virus/Pathogen Mini Kit and Complex200_OBL_V4_DSP protocol (QIAGEN GmbH, Hilden, Germany). Quality control of samples qPCR was performed targeting the Peptidylprolyl Isomerase A (Ppia) housekeeping gene of the mouse as previously described (32). The reaction was performed on LightCycler 480 Platform II using LightCycler® Multiplex RNA Virus Master kit (Roche Lifesciences), and data were analyzed with LightCycler® 480 SW 1.5.1 software using Abs Quant/2nd Derivative Max method.

For *in vitro* MEF experiments, cells were collected at indicated time points and RNA was extracted using NucleoSpin RNA Kit (Macherey Nagel) or QIAamp Viral RNA mini-Kit (Qiagen) on cells and supernatants, respectively, according to manufacturer’s instructions.

Extracted RNA was amplified by real-time (RT) quantitative (q) PCR using Luna Universal One-Step RT-qPCR Kit (NEB) on a StepOnePlus Real-Time PCR System. Pfaffl Model (33) and Bustin MIQE checklist (34) were used for all validations and calculations. To validate our primers, a PCR on a positive control has been performed before dosing 10^-1^ to 10^-12^ 10-fold dilutions of our PCR products using the Denovix DS-11-FX spectrophotometer to evaluate the quantity of DNA in each dilution. Then, the serial dilutions were run in qPCR to validate efficacy, specificity and sensitivity of each couple of primers. Moreover, the exact number of copies in each qPCR reaction was obtained by calculating the “N0 Samples = N0 Standard ∗ Efficacy-ΔCt” before being converted using molecular weight of each targeted gene and Avogadro number. Results obtained were converted to “copies/μg of RNA” for cell lysate samples. Finally, normalization was performed by dividing obtained numbers of RNA copies of the target genes with the deviation of the mouse glyceraldehyde 3-phosphate dehydrogenase (mGAPDH) used as housekeeping gene. Specific set of primers were designed and validated for the detection of mGAPDH (forward: GCATGGCCTTCCGTGTCC; reverse: TGTCATCATACTTGGCAGGTTTCT) and viral NiV-N (forward: GGCAGGATTCTTCGCAACCATC, reverse: GGCTCTTGGGCCAATTTCTCTG).

### Immunofluorescence

MEFs cultured in 12-well plates were washed with PBS 1X and fixed with PFA 4% for 20 min at 4°C. After 2 washes with PBS 1X, cells were permeabilized with TritonX100 0.1%-PBS for 10 min and blocked with of bovine serum albumin (BSA) 2.5%-PBS for 10 min. After 2 washes with PBS-2% FCS-0.1% Tween, cells were incubated 1h at room temperature (RT) with an in-house produced rabbit anti-NiV nucleoprotein (NiV-N) I Ab 1/300 in PBS-2% FCS-0.1% Tween. Plates were then washed 2 times with PBS-2% FCS-0.1% Tween, incubated 1 h at room temperature with a mix of fluorophore-conjugated secondary antibody (anti-mouse AF488, Invitrogen) and DAPI (Merck) 1:1000. Finally, slides were washed 2 times with PBS-2% FCS-0.1% Tween, once with PBS 1X, covered in PBS 1X and stored at 4°C. Fluorescence was evaluated using a Zeiss Axio OBSERVER Z.1 microscope, and photographs were treated using ImageJ software version Java 1.8.0_112.

### Histology and immunohistochemistry

For histopathological evaluation, formalin-fixed specimens were embedded in paraffin wax, 4 μm tissue sections were processed and stained with hematoxylin-eosin-saffron (HES). Immunohistochemistry was performed using the Ventana Discovery Ultra Instrument (Ventana Medical Systems, Roche Diagnostics). The sections were dewaxed and rehydrated. Slides were pre-treated with CC1 (pH 8, Roche Diagnostics) at 98 °C for 40 min. They were then incubated for 30 min at room temperature with blocking buffer (Diagomics), followed by a 1-hour incubation at 37 °C with the primary antibody, rabbit polyclonal anti-NiV-N (1:250). After rinsing, slides were incubated for 30 min at room temperature with a biotinylated goat anti-rabbit IgG antibody (1:200, Dako). Immunoreactive complexes were visualized using streptavidin-conjugated horseradish peroxidase (1:200, Dako) and DAB chromogen (Fisher Scientific). Sections were counterstained with Hematoxylin II and examined using a Nikon Eclipse Ni microscope equipped with a Nikon DS-Ri color camera. Histopathological analysis was performed by a board-certified veterinary pathologist.

## Results

### NiV-Mal replicates at higher levels compared to NiV-Ban in IFNAR KO primary murine cells

The replication and cytopathic effects of NiV-Mal and NiV-Ban were initially analyzed *in vitro* in primary mouse embryonic fibroblasts (MEFs) obtained from WT and IFNAR KO mice sharing the same genetical background employed in further *in vivo* study. Both MEFs genotypes were infected with either NiV strains (MOI = 0.3) and then cell lysates along supernatants were collected at 0h immediately after viral inoculation and at 24h and 48h post infection (Fig.1). NiV-N protein and mRNA levels were analyzed through immunofluorescence (Fig. 1A-1D) and RT-qPCR (Fig. 1E and 1F), respectively. Both cell types were permissive to both NiV strains, although NiV-Ban showed a reduced replication compared to NiV-Mal (Fig. 1A-1F). Also, both viral strains replicated at higher levels in IFNAR KO cells compared to WT cells (Fig. 1A-1D). Moreover, multiple NiV-Mal-induced syncytia appeared 24h post infection and progressively expanded until 48h post infection (Fig. 1A and 1C). Conversely, syncytial multi-nucleated cells were observed only at 48h following NiV-Ban infection (Fig. 1B and 1D) associated with a lower replication rate compared to NiV-Mal infected-cells (Figure 1E and 1F). Globally, NiV-Mal replicated faster than NiV-Ban, resulting in higher levels of viral genomes both produced in the cells and released in the supernatant (Fig. 1A-1F).

**Fig. 1.**
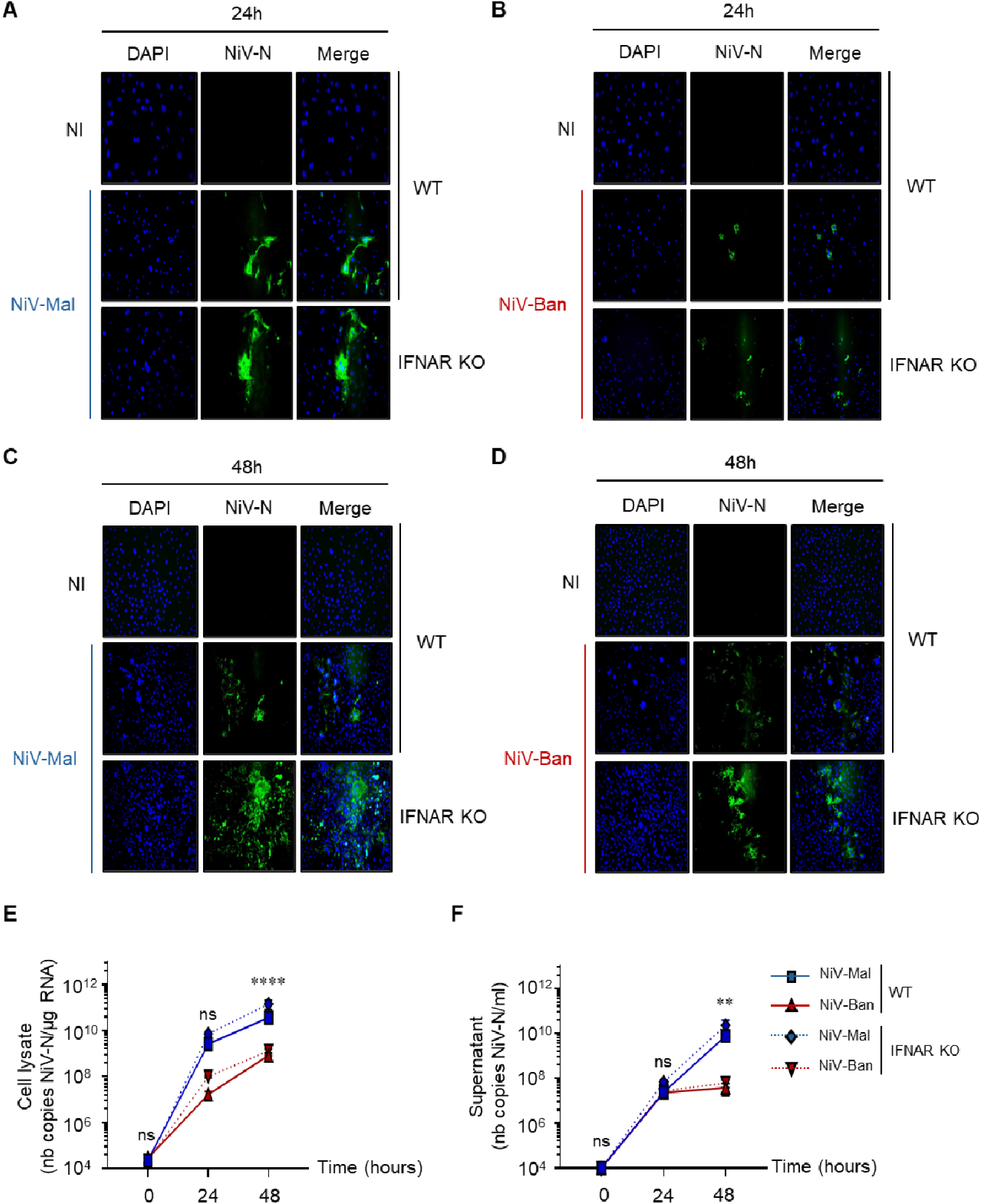
NiV-Mal expands at faster rate than NiV-Ban in primary murine cells. Murine embryonic fibroblasts (MEF) derived from WT or IFNAR KO mice where mock-infected (NI) or infected with NiV-Mal or NiV-Ban at a MOI of 0.3 and cells were cultured for 0h, 24h or 48h (n=3). (A-D) Cells were fixed and stained with DAPI and anti-NiV nucleoprotein (NiV-N) antibody and evaluated by fluorescence microscopy. (E-F) NiV-N mRNA levels from MEF cell lysates and supernatants were assessed by real time RT-qPCR. Data are represented as mean ± standard deviation (SD) of three independent replicates. The difference between NiV-Mal and NiV-Ban in IFNAR KO mice was analyzed using two-way analysis of variance, followed by a Tukey multiple comparison test: ns (not significant); ****p<0.0001.

### NiV-Mal infection is more severe than NiV-Ban following challenge in IFNAR KO mice

We next compared NiV-Mal and NiV-Ban infection in murine models *in vivo.* We infected WT and IFNAR KO mice through IP or IN route with 10^6^ and 10^5^ PFU of virus, respectively of the route of infection, and monitored the animals for 32 days (Fig. 2 and S1).

**Fig. 2.**
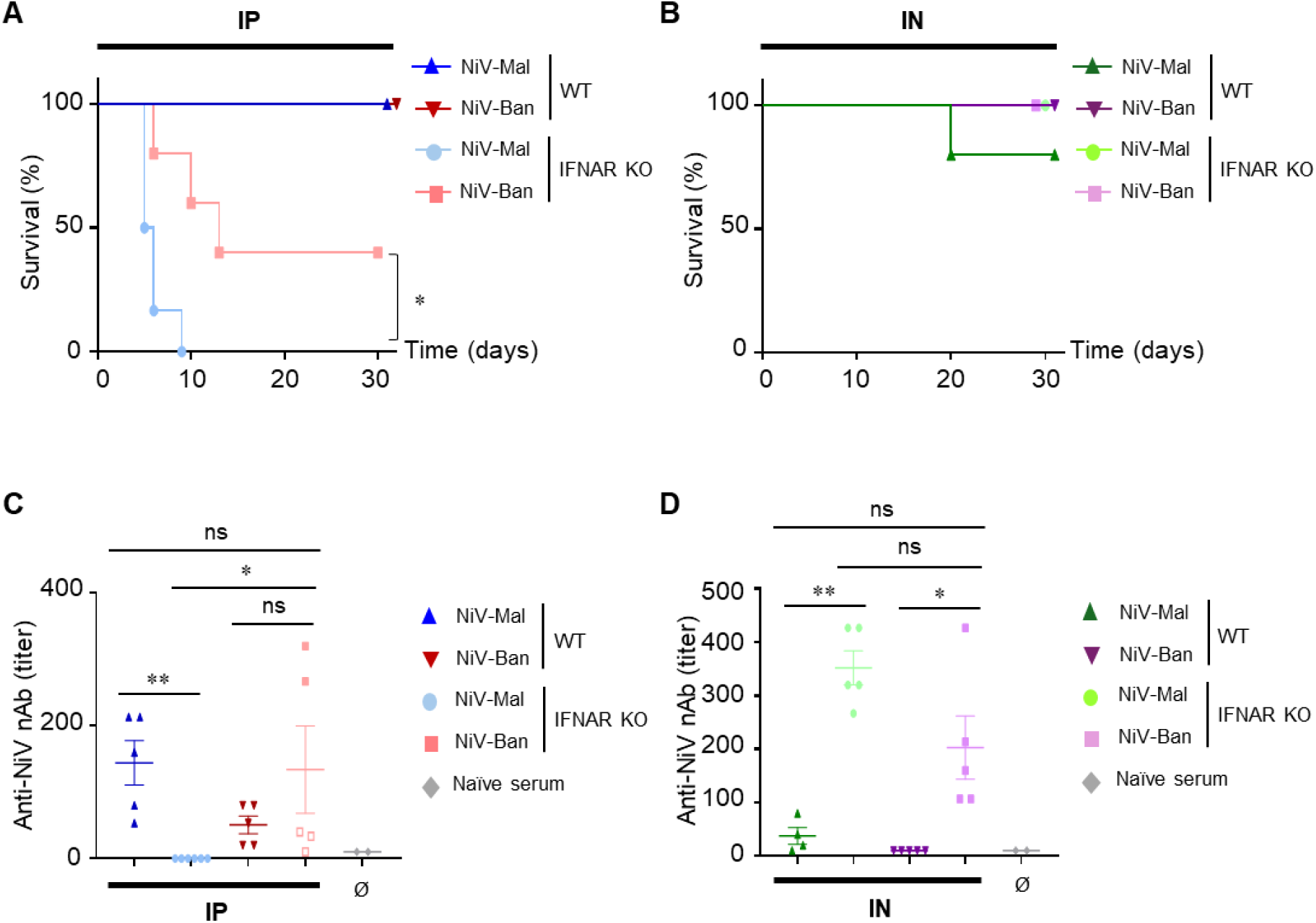
NiV-Mal and NiV-Ban perpetrate different pathogenicity in mice *in vivo*. WT and IFNAR KO mice were infected through an intraperitoneal route (IP) or intranasal route (IN) with 10^6^ or 10^5^ plaque forming units (PFU), respectively, of NiV-Mal or NiV-Ban. All groups were constituted of 5 animals except for NiV-Mal IFNAR KO group, which consisted of 6 mice. (A-B) Survival of infected mice was followed up for 32 days. Survival was analyzed using Gehan-Breslow-Wilcoxon test. *p<0.05. (C-D) Virus-specific neutralizing antibodies (nAb) titer was evaluated by seroneutralization in sera collected at the day of euthanasia (empty dots) or at the end of the experiment (full dots). Serum could not be collected for the NiV-Mal IN-infected mouse which was found dead at d20. Horizontal lines correspond to the average nAb titer of each group. ND = non detected. Data are represented as mean ± standard error mean (SEM). Samples were analyzed using Kruskal Wallis and Dunn’s multiple comparison test, *p<0.05, **p<0.01.

While all WT mice survived IP infections, all NiV-Mal IP-infected IFNAR KO mice succumbed at a mean day of death 6 ± 1.4 days, while NiV-Ban IP infection provoked the death of 60% of mice with a mean day of death of 9.6 ± 2.9 (Fig. 2A). NiV-Mal- and NiV-Ban-infected mice which succumbed to the infection showed similar symptoms, including loss of weight, ruffled hair, facial pain and paralysis (Fig. S1A-S1D). Moreover, 3 IFNAR KO mice IP-infected with NiV-Mal that died at day 5 or day 6 displayed breathing difficulties (Fig. S1C). In contrast, all IN-infected IFNAR KO mice survived until the end of the experiment (Fig. 2B). Intriguingly, one IN-infected WT mouse out of five succumbed NiV-Mal infection and was found dead at d20 (Fig. 2B), despite not showing symptoms throughout the experiment (Fig. S1D). Following IP infection, all WT mice produced specific anti-NiV neutralizing antibodies (nAb) (Fig. 2C). NiV-Mal IP-infected IFNAR KO animals did not produce nAb, consistent with the fact that they all died before day 10, whose period was too short to develop a humoral response (Fig. 2C). Concerning NiV-Ban IP-infected animals, both animals surviving viral challenge produced nAb, thus confirming their initial infection (Fig. 2C). Finally, following IN infection, IFNAR KO mice produced significantly more nAb antibodies than WT mice after infection with either viral strain (Fig. 2D).

### NiV-Mal disseminates at higher levels compared to NiV-Ban following intraperitoneal infection in IFNAR KO mice

To characterize viral replication and expansion, NiV-N RNA levels were quantified in the brain, lung, spleen and liver of infected animals (Fig. 3). As we have previously analyzed NiV-Mal infection in details (25), in the present only one group of NiV-Mal-infected animals was included and monitored during one month (Fig. 3A-3D, left panels). To characterize early events following infection with NiV-Ban strain which has not been tested in IFNAR KO mice before, two groups of mice were infected with NiV-Ban (Fig. 3A-3D, right panels). The first group was monitored until d32, and in the second one single mice were euthanized at defined time points, ie., +4h, d1, d2, d4, d7, d10). For biological accuracy and consistency with our previous study, RT-qPCR data from the two groups of NiV-Ban-infected mice were pooled and represented as “early” (+4h - d1), “mid” (d2 - d7) and “late” (d8 - d32) periods (Fig. 3A-3D, right panels).

**Fig. 3.**
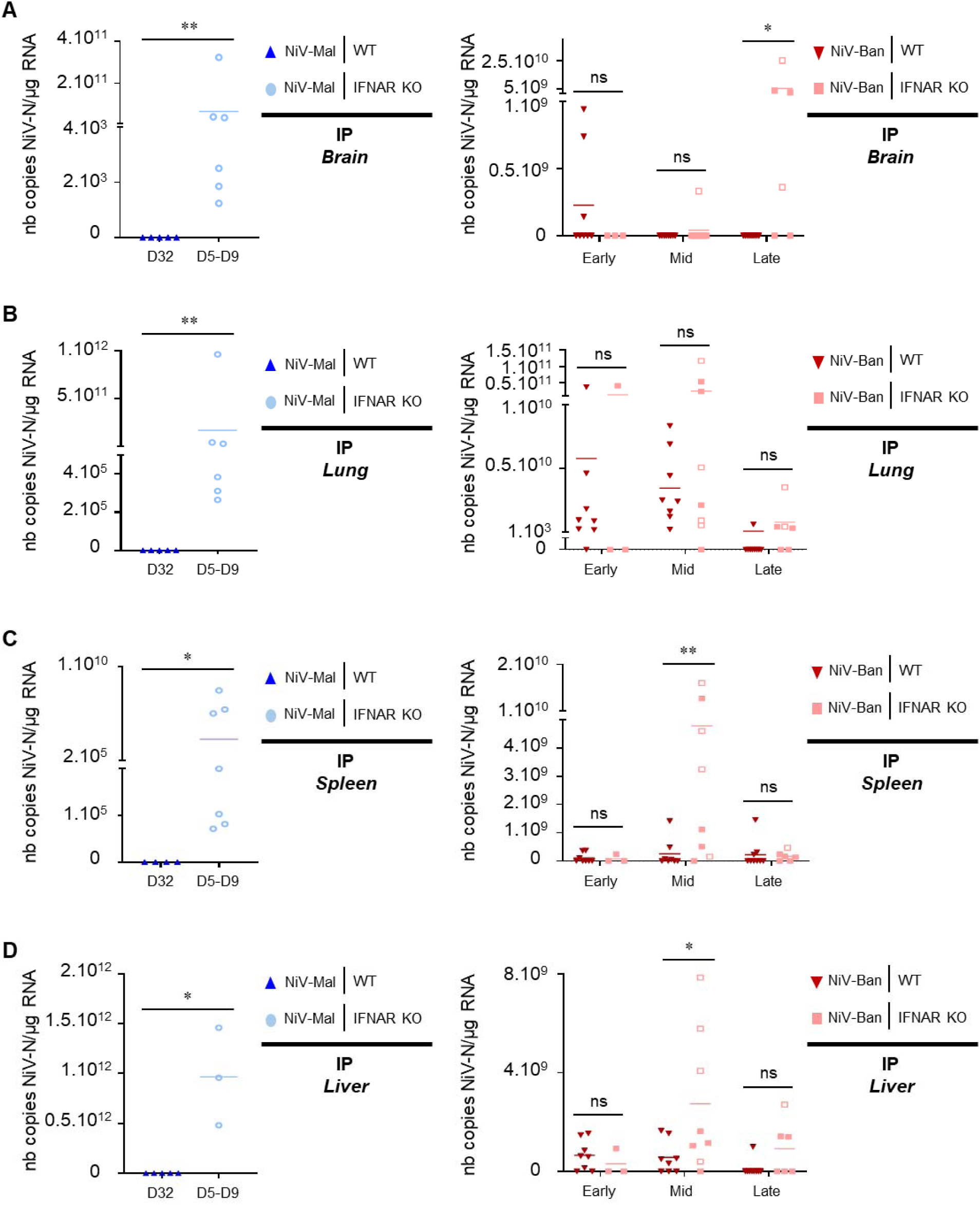
Intraperitoneal NiV infection favors viral dissemination in IFNAR KO mice. (A-D) WT (up blue triangle and down red triangle) and IFNAR KO (light blue circle ang light red square) mice were inoculated intraperitoneally with 10^6^ PFU of NiV-Mal (A-D, left graphs) or NiV-Ban (A-D, right graphs). Following NiV-Mal infection, NiV nucleoprotein (NiV-N) mRNA levels in murine brains (A, left), lungs (B, left), spleens (C, left) and livers (D, left) were harvested either at d32 from surviving WT mice (full dots) or at the day of death of IFNAR KO mice (empty dots) and were assessed by RT-qPCR. Following NiV-Ban infection, NiV-N mRNA levels in murine brains (A, right), lungs (B, right), spleens (C, right) and livers (D, right) were harvested at the day of death (empty dots) or programmed euthanasia (full dots) and were assessed by RT-qPCR. For graphical representation and statistical analyses, mice samples were pooled into early (from d0+4h to d1), mid (from d2 to d7) and late (until d32) phase of infection. Horizontal lines correspond to the average of each group. The statistical difference between IFNAR KO and WT mice groups was analyzed using Mann Whitney test: ns (not significant); *p<0.05; **p<0.01.

Both strains of NiV were detected in the brain (Fig. 3A), lung (Fig. 3B), spleen (Fig. 3C) and liver (Fig. 3D) of IP-infected IFNAR KO mice. No viral RNA was detected in any tested organ of WT animals infected with NiV-Mal analyzed at d32, thus demonstrating effective viral control (Fig. 3A-3D, left panels). NiV-Ban IP-infected WT mice showed active viral replication throughout early, mid and late phases, and still almost completely cleared the infection by d32 (Fig. 3A-3D, right panels). Three WT mice were positive for NiV-Ban in the brain in the early phase of infection, however, viral RNA was no longer detected in the brain throughout the following month (Fig. 3A, right panel). Among IFNAR KO brain samples, 6 out of 6 were virus-positive after NiV-Mal infection, while only 5 out of 17 were virus-positive following NiV-Ban infection (Fig. 3A, left panel and 3A, right panel). Interestingly, 4 out of 6 NiV-Ban-infected IFNAR KO mice had high levels of NiV-N RNA in the brain at the late phase of infection (Fig. 3A, right panel). Out of these 4 mice, one underwent programmed euthanasia at d10, two succumbed due to infection at d10 and d13, and one survived until d32 without clinical signs. While the animals euthanized at d10 and d13 had viral RNA in lung and spleen as well (Fig. 3B and 3C, right panels), the one that reached d32 did not have NiV-N in any other organs except for the brain (Fig. 3A-3D, right panels).

### NiV-Mal and NiV-Ban dissemination is limited following intranasal infection in IFNAR KO mice

Following IN infection with both NiV strains, most animals were negative for NiV-N mRNA, consistent with the fact that, despite one fatal outcome, all other animals survived without major clinical signs (Fig. 4). In contrast, the only WT mouse that succumbed to NiV-Mal infection at d20 was found positive to NiV-N in both brain and lungs (Fig. 4A and 4B, left panels). Contrary to IP-infected animals, no significant differences were detected between IN-infected WT and IFNAR KO mice (Fig. 4A-4D, left and right panels). The most affected organ resulted to be the lung, where virus was detected in approximately 30% of animals (3/10 NiV-Mal-infected mice and 14/45 NiV-Ban-infected mice) (Fig. 4B, left and right panels). Spleens and livers were all negative to NiV-Mal RNA (Fig. 4C and 4D, left panels) and only two IFNAR KO mice were positive to NiV-Ban at mid/late time points, consistent with the IN route of infection (Fig. 4C and 4D, right panels). The brains of two WT and one IFNAR KO animals infected with NiV-Mal were positive for NiV-N RT-qPCR (Fig. 4A, left panel), while the four brain samples that were positive throughout NiV-Ban infection experiment were all collected from IFNAR KO mice (Fig. 4A, right panel).

**Fig. 4.**
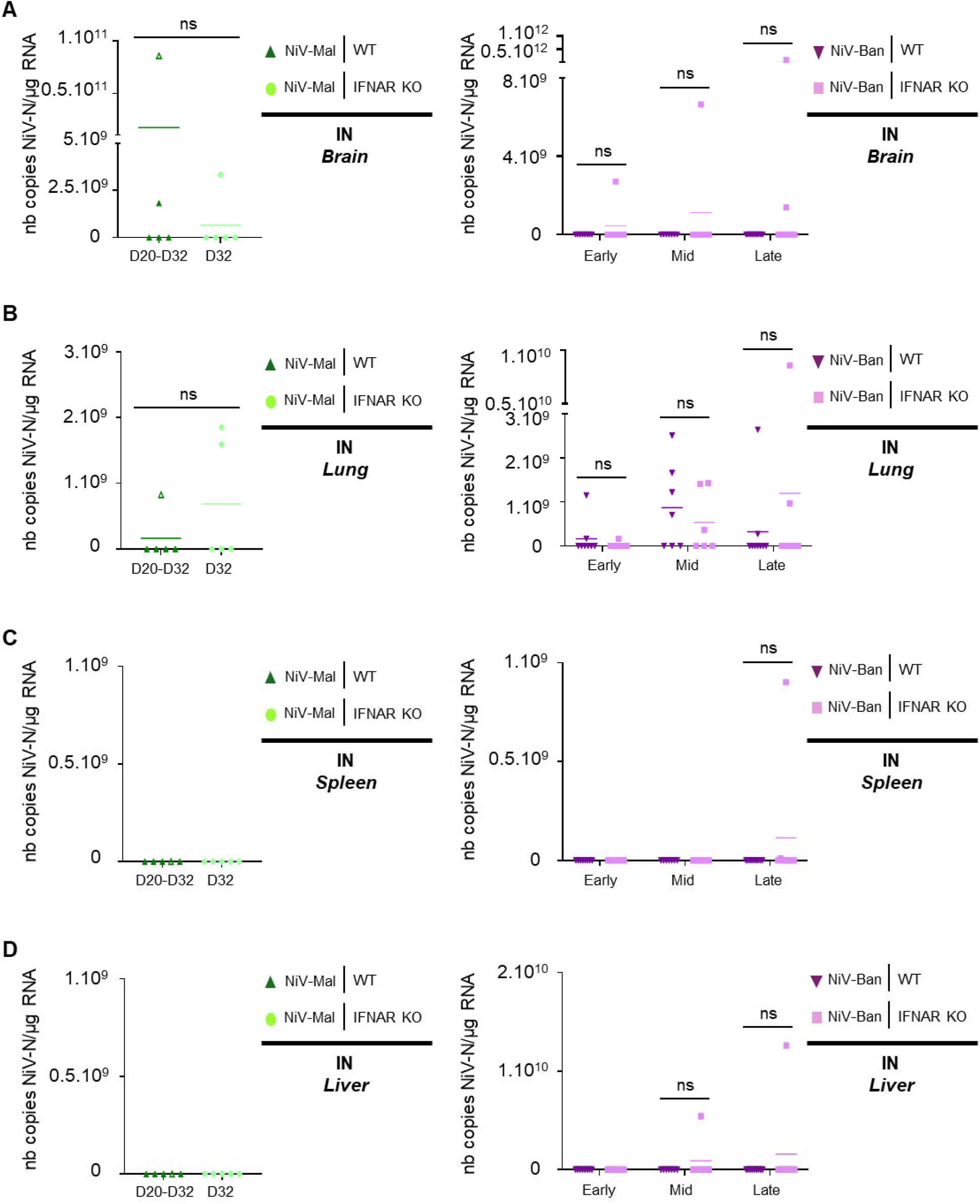
Intranasal NiV infection is controlled in IFNAR KO mice. (A-D) WT (up green triangle and down purple triangle) and IFNAR KO (light green circle ang light purple square) mice were inoculated intraperitoneally with 10^5^ PFU of NiV-Mal (A-D, left graphs) or NiV-Ban (A-D, right graphs). Following NiV-Mal infection, NiV nucleoprotein (NiV-N) mRNA levels in murine brains (A, left), lungs (B, left), spleens (C, left) and livers (D, left) were harvested either at the day of death (d20, empty dot) or d32 from surviving WT mice (full dots) and IFNAR KO mice (full dots) and were assessed by RT-qPCR. Following NiV-Ban infection, NiV-N mRNA levels in murine brains (A, right), lungs (B, right), spleens (C, right) and livers (D, right) were harvested at day 32 (full dots) and were assessed by RT-qPCR. For graphical representation and statistical analyses, mice samples were pooled into early (from d0+4h to d1), mid (from d2 to d7) and late (until d32) phase of infection. Horizontal lines correspond to the average of each group. The statistical difference between IFNAR KO and WT mice groups was analyzed using Mann Whitney test: ns (not significant).

### Intranasal infection of both NiV-Mal and Niv-Ban induces a milder immunopathogenesis compared to intraperitoneal infection in IFNAR KO mice

To better characterize NiV-Mal and NiV-Ban-related pathology, we performed a histopathological analysis and immunolabeling of NiV-N antigen on selected brain, lung and/or spleen sections (Fig. 5, 6, S2 and S3). We prioritized analysis of samples from IP-infected mice that succumbed or were euthanized between d4 and d13 post-infection, corresponding to the critical period of infection (Fig. 5 and 6). Also, NiV-Mal IP-infected WT mice, that all survived until d32, were added to the analysis as controls (Fig. 5, 6, S2 and S3). Indeed, all WT mice resulted negative to NiV-N labeling following both NiV-Mal and NiV-Ban IP infections and did not display major histological signs (Fig. 5A, left panel and S2). Contrary to WT animals, three IFNAR KO mice which succumbed to NiV-Mal infection between d5 and d6 were positive for the presence of NiV-N protein (Fig. 5A, left panel and 5B). NiV-N labeling was found in the brain, in meningeal vessels and olfactory bulb, respectively, with a characterized infiltration of granulocytes (Fig. 5A, left panel and 5B). In the lung, NiV-N localized in bronchial epithelium, some pneumocytes and mononuclear cells of the Bronchi-Associated Lymphoid Tissue (BALT) (Fig. 5B). The detection of some virus in the lung was associated in all 3 animals with vascular lesions including syncytial cells in the endothelium, vascular wall damage, perivascular inflammation and edema (Fig. 6).

**Fig. 5.**
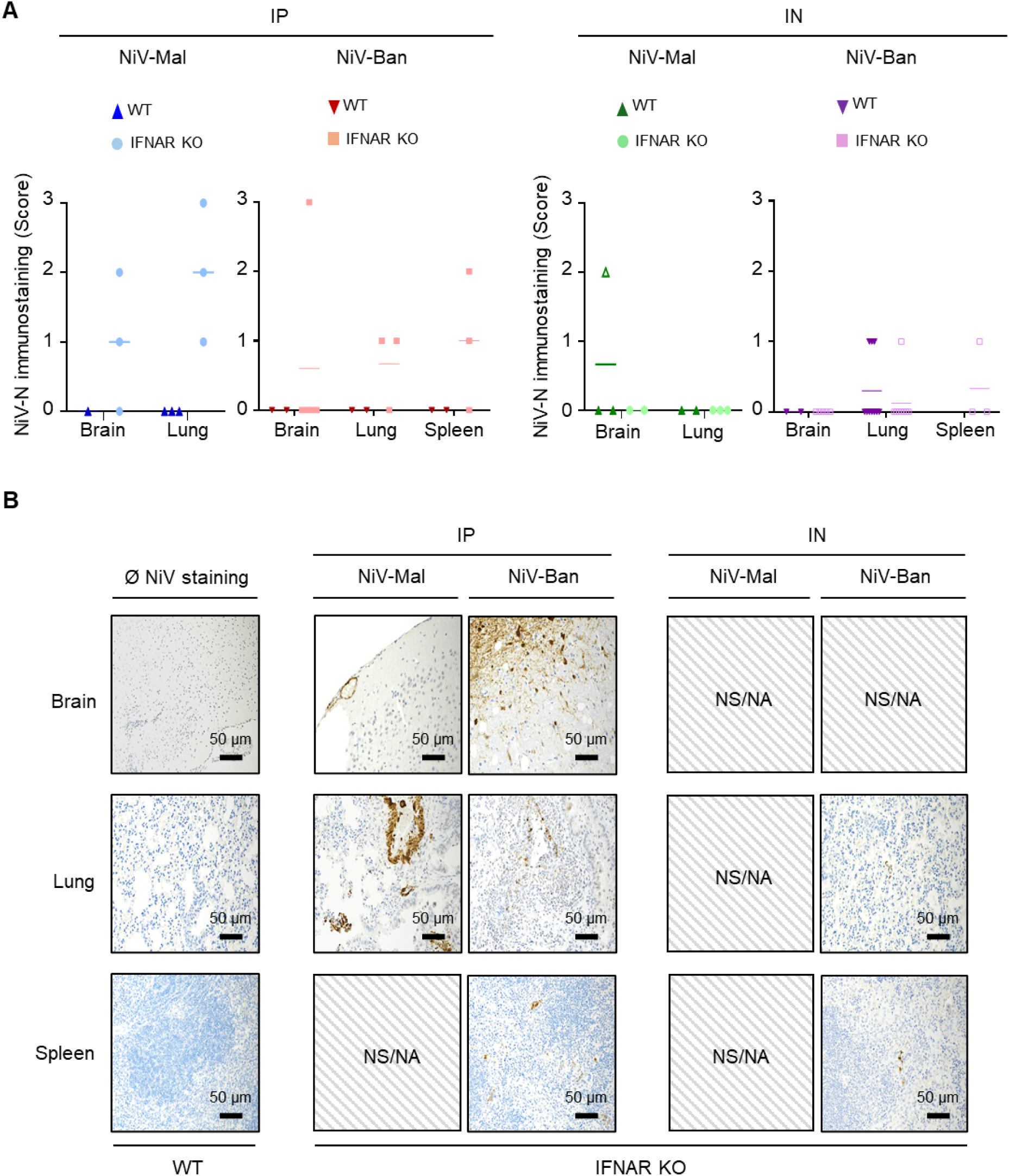
NiV infection level is higher following intraperitoneal infection compared to intranasal infection in IFNAR KO mice. Organs from selected animals were included in paraffin and sections were stained with a rabbit polyclonal anti-NiV-N antibody and DAPI for immunohistochemistry analysis. (A) A scoring system was applied by a veterinary pathologist (0=no labelling; 1=a few positive cells; 2=small clusters of positive cells; 3=multiple large or extensive foci of positive cells) on slides obtained from IP-(left panel) or IN-infected (right panel) organs, respectively. Empty dots represent animals succumbed or euthanized in early/mid late phases of infection (until day 10). (B) Brain, lung and spleens sections from WT mice were used as NiV-negative control conditions while similar sections from IFNAR KO mice were used to characterize the presence of NiV-N antigens following infection. NS/NA represents samples were NiV-N was not shown/not analyzed. Scale bar represents the size of analyzed portions.

**Fig. 6.**
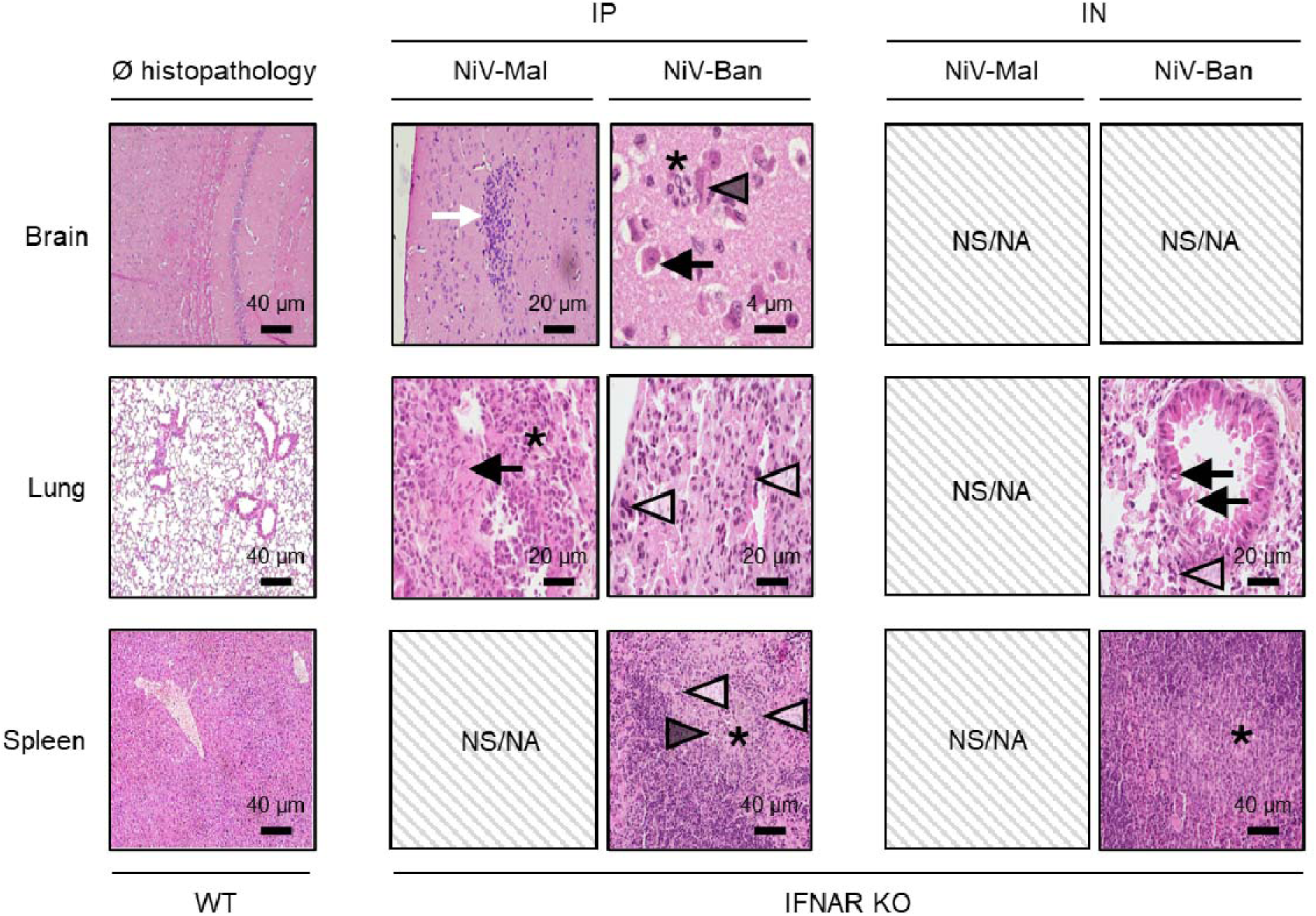
Intraperitoneal NiV infection generates extended immunopathogenesis compared to intranasal infection in IFNAR KO mice. Organs from selected animals underwent hematoxylin-eosin-saffron (HES) staining and were evaluated by a board-certified veterinary pathologist. Brain, lung and spleens sections from WT mice were used as control conditions while similar sections from IFNAR KO mice were used to characterize the presence of immunopathogenic manifestations following NiV infection. White arrow: granulocyte infiltration; Black arrow: inclusion bodies; Black star: hystiocytosis; White arrowhead: syncytial cells; Black arrowhead: necrotic inclusion bodies. NS/NA represents samples were immunopathogenic manifestations were not shown/not analyzed. Scale bar represents the size of analyzed portions.

Regarding NiV-Ban IP infections, three out of four IFNAR KO animals were positive to NiV-N protein immunolabeling (Fig. 5A, left panel). Two of them exerted a positive labeling for NiV antigen in both lung and spleen, with a cellular tropism for pneumocytes and large mononuclear cells of the spleen marginal zone, respectively (Fig. 5B). In the lung, the presence of the virus was associated with perivascular inflammation and syncytial cells, which were diffused throughout alveoli, vessels and bronchi (Fig. 6). In the spleen, syncytia and necrosis were observed mainly in expanded marginal zones that were rich in mononuclear cells, which was indicative of histiocytosis (Fig. 6). In the third IFNAR KO animal, diffused NiV-N staining was found in the brain (lungs and spleens were not available) (Fig. 5A, left panel and 5B). Moreover, this animal displayed extensive lesions corresponding to non-suppurative necrotizing encephalitis and meningitis with perivascular cuffing (Fig. 6).

Regarding the analysis of IN-infected animals, since viral RNA was detected in few samples, we selected animals that were positive for further immunohistochemistry analysis only (Fig. 5A, right panel, 5B, 6, S2 and S3). Out of these 20 animals, only 5 resulted positive for NiV antigen labeling, suggesting that RNA detection is more sensitive than protein labeling (Fig. 5A, right panel and 5B). In both murine genotypes, few cells positive for NiV antigen were detected, and the major histological observation was the presence of perivascular inflammatory cell infiltration in lungs and focal expansion of the marginal zone in spleen (Fig. 6 and S3). The WT mouse that succumbed to NiV-Mal infection at d20 was positive for NiV Ag in the brain and displayed focal encephalitis associated with histiocytosis (Fig. S3). No NiV antigen or histopathological lesions were detected in the brain of other IN-infected animals.

## Discussion

More than 25 years after its first isolation, NiV remains a global health threat due to its high lethality, its human-to-human transmission and the lack of approved therapeutics. Both NiV-Mal and NiV-Ban strains are associated with different symptoms and lethality (35). It is possible that the CFR associated to outbreaks is influenced by multiple factors, such as the viral dose of infection and the route of transmission, as well as socio-economic factors such as Public Health measures and access to healthcare (8,36,37). Indeed, while NiV-Mal emerged only twice in 1998-1999 and 2014, NiV-Ban re-emerged frequently since 2001 (38). Nonetheless, two bats in a city market in Central Java, Indonesia, have been reported positive to NiV-Mal recently, highlighting the risk of spillover from both viral strains (39). The causes responsible for the different diseases associated to NiV-Mal- and NiV-Ban-infected patients remain unclear, and multiple animal models have been tested to characterize its respective pathogenesis (25,28,29,40–47). While non-human primates, hamsters and ferrets succumb to NiV infection, WT mice clear the virus despite being permissive to viral replication (25). However, we previously demonstrated that IFNAR KO mice die following NiV-Mal intraperitoneal challenge due to the lack of type I IFN signaling (25). So far, experiments involving NiV-Ban infection in murine models are lacking and may be of interest to evaluate pathogenic differences between both viral strains *in vivo*. Here, we compared NiV-Mal and NiV-Ban challenge in both WT and IFNAR KO mice infected through the IP or IN route of inoculation.

First, we demonstrated that WT and IFNAR KO primary MEFs are both permissive to NiV-Mal and NiV-Ban infections. Our experiments determined notable differences in the kinetic and the size of formed syncytia in both MEFs genotypes with IFNAR KO cells being more prone to viral dissemination compared to WT cells, along NiV-Mal strain expanding faster than NiV-Ban, thus confirming previous observations made in hamster model (40,42). Then, we investigated such disparities *in vivo* by challenging WT and IFNAR KO mice with each NiV strain through the IP or IN route. Indeed, consistent with our *in vitro* observations, higher lethality was observed in IFNAR KO following NiV-Mal infection compared to NiV-Ban infection through the IP route, while all WT mice survived. Surprisingly, all IFNAR KO mice including animals infected with NiV-Ban, known to be associated with respiratory distress syndrome in humans, survived IN infection. Indeed, contrary to the IP route, IN inoculation led to a subclinical infection with both NiV strains in IFNAR KO mice, thus confirming previous observations where the IN route lead to an attenuated disease (25). Also, the lower inoculation dose (10^5^ PFU) used for IN challenge in this study compared to the dose used in our previous report (10^6^ PFU) might be responsible for the absence of fatal outcomes (25). Overall, contrary to the human-associated NiV disease observed in African green monkeys, rodents including our murine models may fail to recapitulate NiV morbidities and outcomes (40,46), potentially due to species-specific factors, such as a more developed basal immune environment in the respiratory tract when compared with primates (48–50).

Concomitantly, IN-inoculated and surviving IP-inoculated IFNAR KO mice produced significantly higher nAb titers compared to WT mice following NiV infection with both strains. This demonstrates that despite the lack of IFN-I signaling-dependent activity and higher viral replication, the adaptive immune response was strong enough to control viral growth. Interestingly, the lack of visible pathogenic signs in both WT and IFNAR KO mice following IN infection may be related to a major role of type III IFNs in the defense against pathogens in mucosal barriers such as the respiratory tract, as previously described (51–53). Regarding brain infection, few analyzed samples were positive following IN inoculation, suggesting differences in viral entry in the central nervous system (CNS) through the olfactory bulb as described in hamsters (54). Intriguingly, in the mouse model, the hematogenous route resulted in a higher propension of the virus to reach the CNS, as observed in IP-infected mice. Globally, considering the absence of clinical symptoms and scarce viral propagation, murine models do not appear to be very susceptible to IN infection. However, further tests with a higher inoculation dose or with combined KOs could clarify differences between NiV-Mal and NiV-Ban *in vivo* following IN inoculation.

Interestingly, a different cell tropism was observed in the lungs between the two viral strains following IP infection. NiV-Mal-associated antigen was mainly detected in the bronchi and BALT, while NiV-Ban mostly infected alveolar pneumocytes. Moreover, syncytia in the lungs were centered on blood vessels with NiV-Mal but were diffused throughout lung parenchyma and bronchi with NiV-Ban. Further investigations are necessary to understand whether the observed cellular tropism and syncytia distribution are host-specific and whether they have consequences on viral shedding. Furthermore, virus-induced cell-to-cell fusion has been demonstrated to be associated with STING responses and may contribute to disrupt the blood-brain barrier following NiV infection as well, but syncytia formation has not always been thoroughly analyzed in animal experiments in the literature (27,55–57). Thus, systematic reporting of syncytia formation in future studies could contribute to shed light into differences between NiV strains.

Concerning the spleen, two IFNAR KO animals which succumbed to NiV-Ban IP infection displayed foci of NiV-N antigen positive large mononuclear cells, corresponding to macrophages, in the mantle zone. This demonstrates that despite antigen interaction/potential internalization by antigen-presenting-cells (APCs) in the spleen, the immune response was not fast and/or strong enough to control infection.

In the brain, no strain-specific differences in cell tropism were observed and viruses were found in diverse compartments, including meninges and olfactory bulb. This corroborates the observation that NiV can potentially distribute and induce lesions throughout the whole brain, as recently reported by Goldin *et al*. following NiV-Ban infection in AGMs (58). Moreover, while both viral strains could induce similar lesions in the brain, the capacity of NiV-Ban to reach the brain in humans is restrained due to the fast and severe respiratory syndrome it induces, thus limiting its ability to further affect the brain. This was supported by Goldin *et al.*, who reported that treatments counteracting NiV-Ban respiratory disease ultimately led to the death of monkeys through encephalitis (58).

Altogether, we tested IFNAR KO mice as potential experimental model for both NiV-Mal and NiV-Ban infections. IP infection resulted in lethal outcome in 100% of NiV-Mal- and 60% of NiV-Ban-infected mice in this model with viral replication detected in all tested organs. Conversely, IN inoculation induced a subclinical infection with both viral strains. While hamsters and AGMs are the most suitable models to study NiV infection due to their natural susceptibility to the virus, mice could represent a useful alternative due to the large number of immune reagents, genetic tools and global knowledge in murine immune system available. Interestingly, the NiV-Ban IP-infected IFNAR KO model is of high interest to study chronic NiV infection as NiV-positive animals without symptoms were identified after 32 days, thus representing a potential model to challenge therapeutics against cerebral infection. Moreover, the possibility to produce specific KO mice may allow further dissect the contribution of specific immune factors involved in the control of NiV. In conclusion, mice represent a valid small animal model for specific immunopathogenic and fundamental investigations in the study of NiV infection.

## CRediT authorship contribution statement

Conceptualization: MI, BH; Acquisition and analysis of data: LA, BP, DD, IR, JSp, JSk, TL, OT; Writing Original Draft: LA, BH, OR, MI; Providing critical methodology: UK, JSp, JSk; Supervision: BH, MI; Funding Acquisition: BH, MI.

## Supporting information

Supporting informations

## Acknowledgements

The study was supported by ERINHA-Advance (ICONIM) and by Aviesan Sino-French agreement on Nipah virus study and by INSERM. We are grateful to all the members of the group Immunobiology of viral infection at CIRI, and Audrey Richard and Claudia Filippone from ERINHA, for the help in the realization of this study. We thank Professor Yuke-Fun Chan of University of Malaya for kindly providing Nipah virus Malaysia strain. We thank to Center for Disease Control and Prevention, Atlanta, USA, for kindly providing Nipah virus Bangladesh isolate. We acknowledge the contribution of the SFR Biosciences (UMS3444/CNRS, US8/Inserm) in Lyon.

## Supporting information

**S1 Fig. Different pathogenic signatures of NiV Mal and NiV-Ban infection in mice *in vivo*.** WT and IFNAR KO mice were infected through an intraperitoneal route (A, C) or intranasal route (B, D) with 10^6^ or 10^5^ PFU, respectively, of NiV-Mal or NiV-Ban. All groups were constituted of 5 animals except for NiV-Mal IFNAR KO group, which consisted of 6 mice. Animals were monitored for 32 days for weight (A-B) and clinical signs with established clinical scores (C-D).

**S2 Fig. NiV infection is attenuated in WT mice.** Organs from selected animals were included in paraffin and sections were stained with a rabbit polyclonal anti-NiV-N antibody and DAPI for immunohistochemistry analysis. (A) Brain, lung and spleens sections from WT mice were used as NiV-negative control conditions while similar sections from WT mice sensitive to NiV infection were used to characterize the presence of NiV-N antigens. NS/NA represents samples were NiV-N was not shown/not analyzed. Scale bar represents the size of analyzed portions.

**S3 Fig. Intraperitoneal and intranasal NiV infections generate mild immunopathogenesis in WT mice.** Organs from selected animals underwent hematoxylin-eosin-saffron (HES) staining and were evaluated by a board-certified veterinary pathologist. Brain, lung and spleens sections from WT mice were used as control conditions while similar sections from WT mice sensitive to NiV infection were used to characterize the presence of immunopathogenic manifestations. Black star: hystiocytosis; White arrowhead: syncytial cells. NS/NA represents samples were immunopathogenic manifestations were not shown/not analyzed. Scale bar represents the size of analyzed portions.

