## Supplementary figures and images for "Nipah virus Malaysia and Bangladesh strain-induced pathogenesis in mice lacking type I interferon receptor signaling"

### Supporting informations

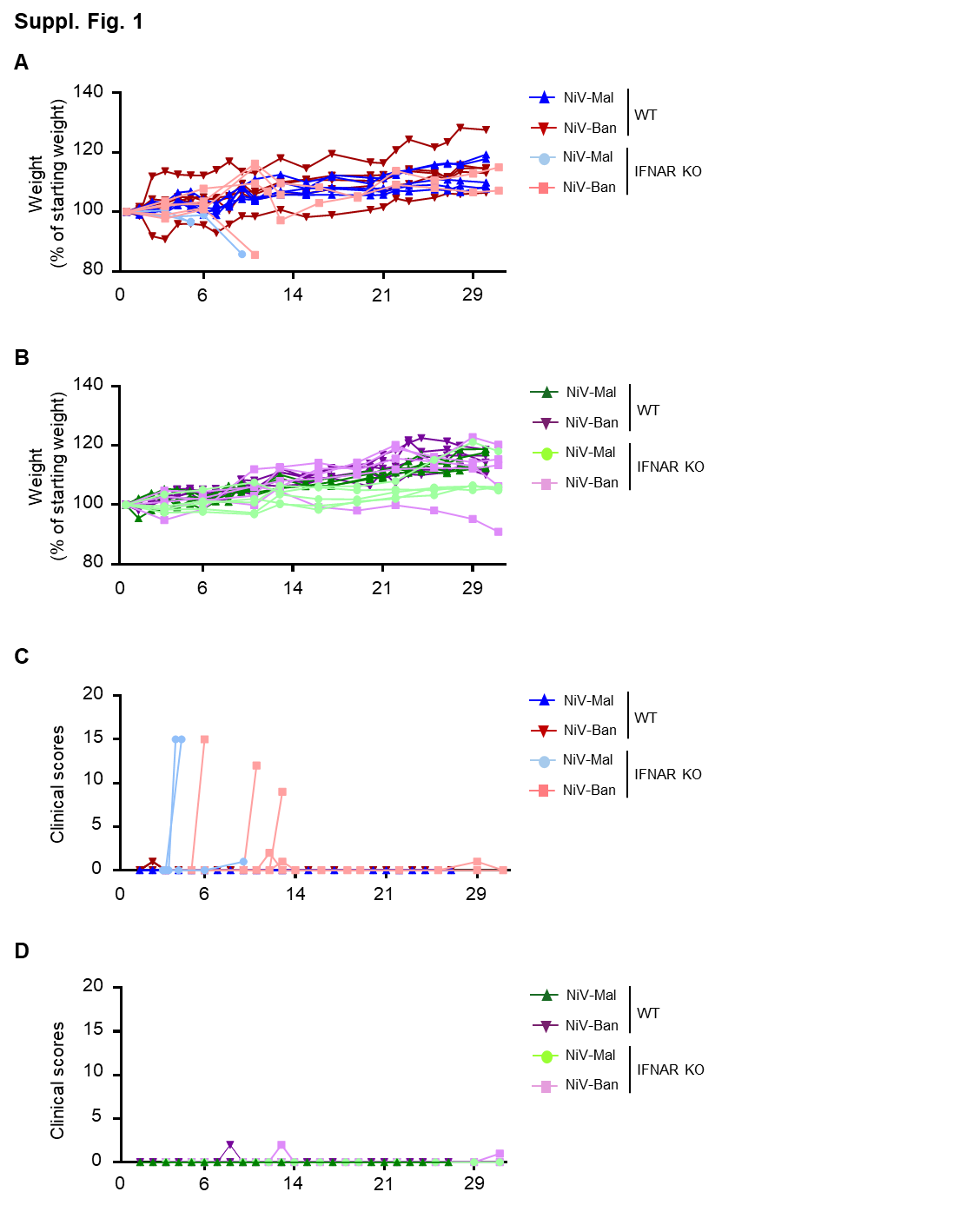

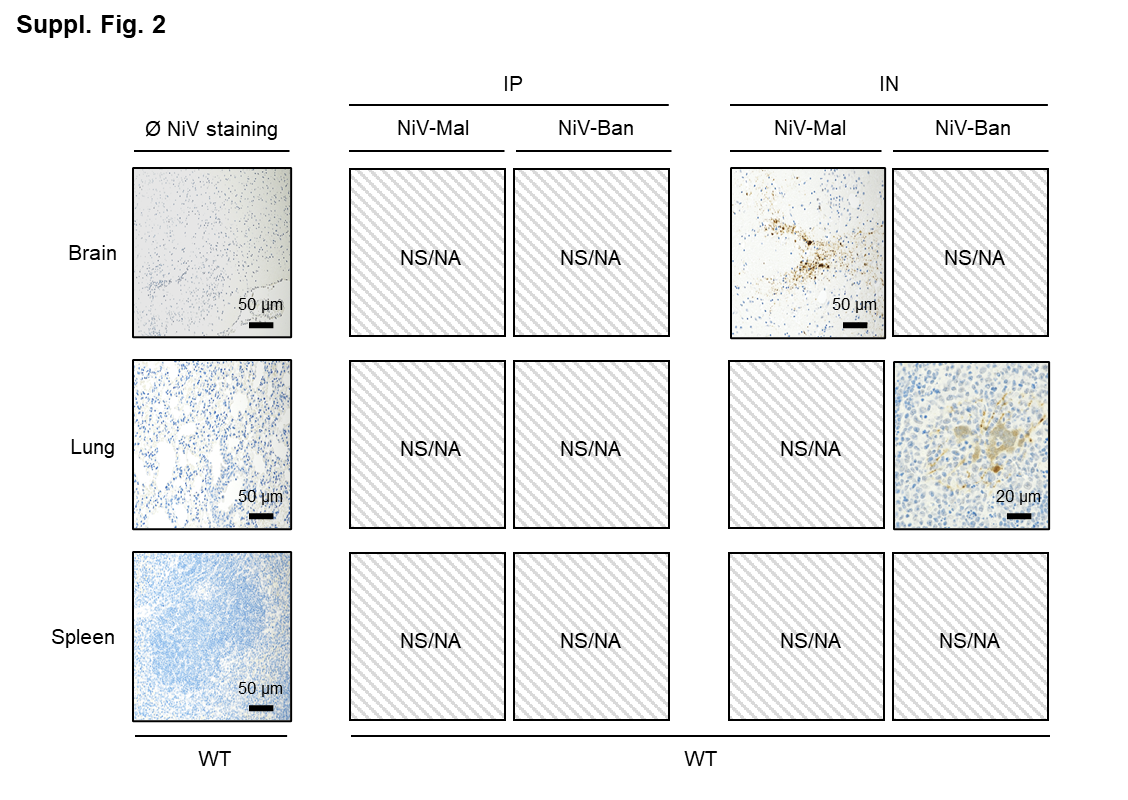

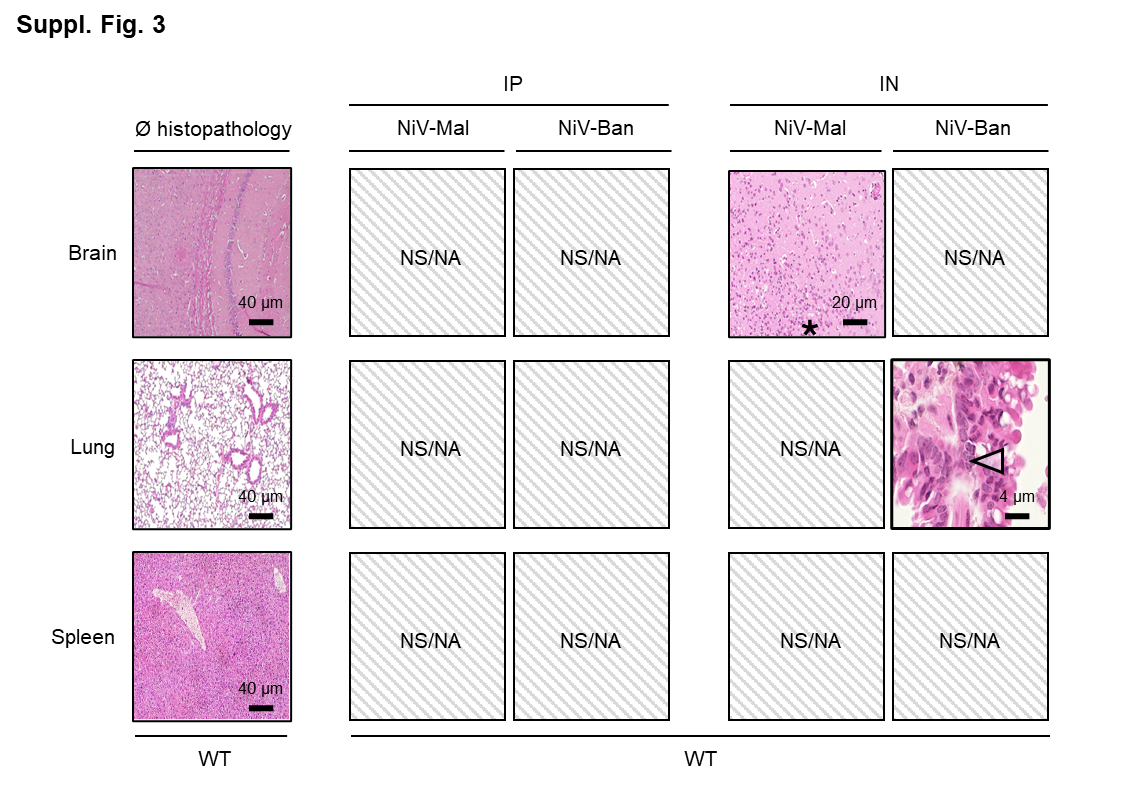
